# Reproducible, high-dimensional imaging in archival human tissue by Multiplexed Ion Beam Imaging by Time-of-Flight (MIBI-TOF)

**DOI:** 10.1101/2021.10.14.464455

**Authors:** Candace C. Liu, Marc Bosse, Alex Kong, Adam Kagel, Robert Kinders, Stephen M. Hewitt, Sean C. Bendall, Michael Angelo

## Abstract

Multiplexed ion beam imaging by time-of-flight (MIBI-TOF) is a form of mass spectrometry imaging that uses metal labeled antibodies and secondary ion mass spectrometry to image dozens of proteins simultaneously in the same tissue section. Working with the National Cancer Institute’s (NCI) Cancer Immune Monitoring and Analysis Centers (CIMAC), we undertook a validation study, assessing concordance across a dozen serial sections of a tissue microarray of 21 samples that were independently processed and imaged by MIBI-TOF or single-plex immunohistochemistry (IHC) over 12 days. Pixel-level features were highly concordant across all 16 targets assessed in both staining intensity (*R*^*2*^ = 0.94 ± 0.04) and frequency (*R*^*2*^ = 0.95 ± 0.04). Comparison to digitized, single-plex IHC on adjacent serial sections revealed similar concordance (*R*^*2*^ = 0.85 ± 0.08) as well. Lastly, automated segmentation and clustering of eight cell populations found that cell frequencies between replicates yielded an average correlation of *R*^*2*^ = 0.92 ± 0.06. Taken together, we demonstrate that MIBI-TOF, with well-vetted reagents and automated analysis, can generate consistent and quantitative annotations of clinically relevant cell states in archival human tissue, and more broadly, present a scalable framework for benchmarking multiplexed IHC approaches.

## Introduction

Immunohistochemistry (IHC) is commonly used in clinical diagnostics and basic research to visualize proteins in intact tissue using chromogenic or fluorescent reporters.^1–3^ IHC staining is used routinely to guide diagnoses and therapeutic selection in the vast majority of solid tissue malignancies.^4–6^ Although it remains an indispensable tool in anatomic pathology, chromogenic IHC has inherent limitations that hinder quantitative interpretation and prevent routine multiplexed staining.^1,7 8–10^ These drawbacks are particularly limiting in the field of cancer immunotherapy, where accurate evaluation of the tumor immune microenvironment requires the simultaneous mapping of dozens of proteins.^11,12^ The emerging field of spatial-omics with a multitude of analytical solutions is working towards replacement of conventional IHC-driven decision making.^11,13^ While it is clear that new technologies and assays have the potential to generate new types of data, it is unclear whether they can reliably replicate the decision-making information in current ‘gold standard’ assays.

Our lab has developed multiplexed ion beam imaging by time-of-flight (MIBI-TOF), which avoids the limitations of optical imaging by using secondary mass spectrometry to image dozens of proteins on the same tissue section.^14,15^ In the place of chromogenic or fluorescent reporters, MIBI-TOF uses primary antibodies that are labeled with isotopically enriched metal reporters that can be cleanly delineated and quantified using time-of-flight (TOF) mass spectrometry. Tissue sections are treated with all metal labeled primary antibodies simultaneously using a simple protocol that does not include secondary antibodies, enzymatic amplification, or cyclical staining. During MIBI-TOF analysis, the tissue is sputtered by a primary ion beam in a pixel-by-pixel fashion that liberates the metal tags as secondary ions that are subsequently quantified by TOF. Our lab routinely quantifies 40 targets simultaneously and are currently working towards increasing this capability to 60 or more.^16–18^ Notably, MIBI-TOF is compatible with formalin-fixed, paraffin-embedded (FFPE) samples and can detect both low and high abundant targets with a dynamic range that spans six orders of magnitude.^15 19–21^

For technologies like MIBI-TOF to be used in large translational studies and ultimately for routine clinical diagnostics, robustness and reproducibility studies are needed, as are standardized workflows for interpreting these complex data. Furthermore, it is important to show that new technologies are concordant with existing, established clinical assays. In collaboration with the National Cancer Institute (NCI) as part of the Cancer Immune Monitoring and Analysis Centers-Cancer Immunologic Data Commons (CIMAC-CIDC) network^22^, we benchmarked the reproducibility of multiplexed antibody staining using MIBI-TOF. To achieve this, we compared MIBI-TOF imaging data across six independent replicates of adjacent tissue microarray (TMA) serial sections. Antibody staining and MIBI-TOF imaging of each serial section were carried out independently and randomized with respect to all experimental parameters. Additionally, we assessed MIBI-TOF concordance with single-plex IHC chromogenic staining. In keeping with the goal of CIMAC-CIDC to develop comprehensive, standardized immune monitoring analysis for biomarker discovery, we show that MIBI-TOF is a highly reproducible assay and concordant with single-plex IHC.

## Materials and Methods

### Tissue microarray construction and sectioning

A tissue microarray (TMA) was constructed using human FFPE blocks from Stanford Pathology. The TMA consisted of disease-free controls as well as multiple types of carcinomas, sarcomas, and central nervous system lesions. A table of the cores included in this study can be found in Table 1 (1 mm cores). For each TMA tissue block, 13 consecutive serial sections (4 μm section thickness) were acquired. The IHC and MIBI-TOF recuts were alternated so that for the IHC concordance analysis, MIBI-TOF and IHC could be compared in adjacent sections. The assays performed on each section were as follows:

**Table 1:** Description of tissue cores and parameter randomization.

| Slide Number | Replicate 1 | Replicate 2 | Replicate 3 | Replicate 4 | Replicate 5 | Replicate 6 |
| --- | --- | --- | --- | --- | --- | --- |
| Stain Order | 1 | 2 | 6 | 5 | 3 | 4 |
| Day of Analysis | 5 | 2 | 6 | 3 | 4 | 1 |

**Table 1: Description of tissue cores and parameter randomization**
| FOV | Order of Acquisition |  |  |  |  |  |
| --- | --- | --- | --- | --- | --- | --- |
| CONTROL TONSIL | 14 | 4 | 17 | 1 | 19 | 1 |
| CONTROL TONSIL | 18 | 20 | 21 | 7 | 20 | 2 |
| CONTROL TONSIL | 19 | 10 | 6 | 5 | 13 | 3 |
| CONTROL TONSIL | 11 | 1 | 2 | 18 | 5 | 4 |
| CONTROL LYMPH NODE | 9 | 3 | 18 | 2 | 11 | 5 |
| CONTROL LYMPH NODE | 15 | 6 | 11 | 12 | 10 | 6 |
| CONTROL LYMPH NODE | 3 | 7 | 15 | 9 | 4 | 7 |
| CONTROL THYMUS | 7 | 11 | 14 | 20 | 3 | 8 |
| CONTROL THYMUS | 5 | 12 | 5 | 13 | 8 | 9 |
| CONTROL SPLEEN | 17 | 19 | 16 | 16 | 21 | 10 |
| SALIVARY WARTHIN SALIVARY | 2 | 17 | 4 | 4 | 9 | 11 |
| HIGH GRADE, CARCINOMA BLADDER | 16 | 15 | 12 | 19 | 18 | 12 |
| CONTROL PLACENTA | 8 | 9 | 20 | 14 | 7 | 13 |
| DUCTAL CARCINOMA BREAST | 20 | 18 | 3 | 21 | 1 | 14 |
| DUCTAL CARCINOMA BREAST | 1 | 2 | 13 | 15 | 12 | 15 |
| LEIOMYOSARCOMA, MET SOFT TISSUE | 12 | 13 | 7 | 8 | 6 | 16 |
| HIGH-GRADE MYXOFIBROSARCOMA SOFT TISSUE | 6 | 16 | 19 | 10 | 15 | 17 |
| FOREIGN BODY GIANT CELLS SOFT TISSUE | 21 | 21 | 9 | 6 | 14 | 18 |
| ADENOCARCINOMA COLON | 10 | 8 | 1 | 3 | 2 | 19 |
| PLEOMORPHIC DERMAL SARCOMA SKIN | 13 | 14 | 10 | 11 | 16 | 20 |
| SQUAMOUS CELL CARCINOMA SKIN | 4 | 5 | 8 | 17 | 17 | 21 |

Section 1: H&E

Sections 2,4,6,8,10: Chromogenic IHC

Sections 3,5,7,9,11,13: MIBI-TOF full panel staining

Section 12: MIBI-TOF unstained control

### Antibody preparation

The full MIBI-TOF panel containing 16 antibodies can be found in Table 2. The lyophilized antibody panel was obtained from Ionpath (“Cell Classification” human panel). Individual metal labeled antibodies for IHC were also obtained from Ionpath: CD3 (D7A6E), CD8 (C8/144B), CD68 (D4B9C), Pax5 (D7H5X), PanCK (AE1/AE3). Prior to staining, the lyophilized antibody panel was suspended in antibody diluent buffer (TBS-IHC tween, Donkey serum 3%) and filtered with a 0.1 µm filter (Millipore).

**Table 2:** MIBI-TOF staining panel. Targets used for IHC concordance analysis are highlighted in green.

| Target | Clone |
| --- | --- |
| dsDNA | 35I9 DNA |
| beta-tubulin | D3U1W |
| CD4 | EPR6855 |
| CD11c | EP1347Y |
| CD56 | MRQ-42 |
| CD31 | EP3095 |
| CD68 | D4B9C |
| CD8 | C8/144B |
| CD3 | D7A6E |
| Vimentin | D21H3 |
| CD20 | L26 |
| HLA DR | EPR3692 |
| CD45 | 2B11 & PD7/26 |
| Na-K-ATPase alpha 1 | D4Y7E |
| Pax5 | D7H5X |
| PanCK | AE1/AE3 |

### Tissue staining

Tissue staining for the six MIBI-TOF recuts was carried out over six separate days, randomized with respect to serial section number (Table 1). Detailed protocols can be found here: dx.doi.org/10.17504/protocols.io.byzrpx56, dx.doi.org/10.17504/protocols.io.bf6ajrae, dx.doi.org/10.17504/protocols.io.bhmej43e. We used the Sequenza manual staining system for both MIBI-TOF and IHC staining (dx.doi.org/10.17504/protocols.io.bmc6k2ze).

Briefly, slides were baked at 70°C overnight followed by deparaffinization and rehydration with successive washes of 30 seconds and 3 dips each in xylene (3 washes), 100% ethanol (2 washes), 95% ethanol (2 washes), 80% ethanol (1 wash), 70% ethanol (1 wash), and ddH2O (2 washes) with a Leica ST4020 Linear Stainer (Leica Biosystems). Tissues next underwent antigen retrieval by submerging slides in 3-in-1 Target Retrieval Solution (pH 9, DAKO Agilent) and incubating at 97°C for 40 minutes in a Lab Vision PT Module (Thermo Fisher Scientific). After cooling to room temperature, slides were assembled with a cover plate on a Sequenza rack and washed with PBS wash buffer. The tissue was blocked for 1 hour at room temperature with 1x TBS IHC Wash Buffer with Tween 20 with 3% (v/v) normal donkey serum (Sigma-Aldrich), 0.1% (v/v) cold fish skin gelatin (Sigma Aldrich), 0.1% (v/v) Triton X-100, and 0.05% (v/v) sodium azide. The blocking buffer was washed by adding 200 μL of antibody diluent buffer in the Sequenza upper chamber. After the diluent buffer flow through, 120 μL of the suspended panel of antibodies was added to the slides. The Sequenza rack was then placed at 4°C overnight (16 hr).

Following the overnight incubation, slides were washed twice with 1 mL of PBS wash buffer and fixed in a solution of 2% glutaraldehyde (Electron Microscopy Sciences) solution in low-barium PBS for 5 minutes. Slides were successively washed with 30 seconds and 3 dips per wash in PBS (1 wash), 0.1 M Tris at pH 8.5 (3 washes), ddH2O (2 washes), and then dehydrated by washing in 70% ethanol (1 wash), 80% ethanol (1 wash), 95% ethanol (2 washes), and 100% ethanol (2 washes). Slides were dried under vacuum prior to imaging.

Single-plex chromogenic IHC was performed for five targets using the same antibody clone as used in the MIBI-TOF staining panel (Table 2). Dewaxing, epitope retrieval, blocking, hybridization, and washing for IHC tissue sections were identical to that of MIBI, with the addition of blocking endogenous peroxidase activity with 3% H_2_O_2_ (Sigma Aldrich) in ddH2O after epitope retrieval. After overnight primary antibody staining, tissues were washed twice with 1 mL wash buffer and the antigen:antibody complex detected with the ImmPRESS universal (Anti-Mouse/Anti-Rabbit) kit (Vector labs). Chromogenic solution (DAB) was incubated for 40 s and the reaction was stopped with PBS. The slides were counterstained with hematoxylin.

H&E tissue sections were reviewed to identify missing tissue cores, cores with large areas of necrosis, tissue folding, and any other macroscopic defect. All IHC and MIBI-TOF data were manually reviewed for excessive background staining.

### MIBI-TOF imaging

The order of image acquisition for each recut was randomized with respect to section number and carried out on six separate days (Table 1). Within each run for each recut, the order of acquisition for the TMA cores was randomized as well (Table 1). Area normalized Xe+ primary ion dose of 9nA*hr*mm^-2^ was used for all MIBI-TOF image acquisitions. At the end of each imaging run, sensitivity and mass resolution were quantified using a molybdenum foil standard (Supplementary Figure 1).

### Image processing

MIBI-TOF data processing was performed using a standardized pipeline from Ionpath. During data extraction, to only capture the monoatomic peaks that correspond to the monoatomic isotopic labels and avoid the polyatomic ions containing hydrogen, we integrated all peaks using a mass window of (X-0.3, X) where X is the nominal mass-to-charge (m/z) for each metal reporter tag.

Background signal arising from bare slide was removed as described previously using predefined thresholds for Ba, Ta, and Au.^16,23^ Isobaric interferences arising from diatomic reporter adducts and polyatomic hydrocarbons were removed iteratively using empirically defined compensation coefficients (Supplementary Table 1). Lastly, we used a combination of several density metrics, including reachability density, connectivity, and Voronoi tessellation to estimate and remove noise from each channel (Supplementary Table 2). These density metrics leverage both spatial and intensity information.

Hematoxylin counterstained IHC slides were digitized with the NanoZoomer slide scanner at 40x magnification (Hamamatsu). The threshold for hematoxylin and each DAB channel was set collectively for all images in the staining batch using a DAB channel extracted from the original counterstained image via spectral unmixing. An initial threshold approximation was set with automatic image histogram thresholding. The thresholds were then confirmed by three trained users on controls and additional images for validation and adjusted until consensus was reached (Supplementary Table 3).

### Marker quantitation

In all MIBI-TOF images, for each marker, the frequency and mean intensity of positive pixels in each tissue core was quantified. The resultant value for percent positive pixels (PPP) and mean pixel intensity (MPI) for these images was then normalized with respect to the signal intensity of endogenous carbon-containing mass peaks originating from the tissue itself. Peaks at 36, 37 and 38 m/z corresponding to 3C, 3C1H, and 3C2H were integrated for each image (referred to here as *C*_*i*_) and the average of these values across all images in all six TMAs was calculated (referred here to *C*_*avg*_). To calculate the normalization coefficient (*C*_*N*_) for a given tissue core, *C*_*i*_ for that tissue core was divided by *C*_*avg*_ (*C*_*N*_ = *C*_*i*_ / *C*_*avg*_). Median and standard deviation for *C*_*N*_ across all tissue cores was 1.03 ± 0.3. For IHC images, the frequency of positive pixels was quantified using deconvolved and thresholded DAB images (PPP).

### Reproducibility and concordance

To assess replicate reproducibility, least squares linear regression was used to compare PPP for tissue cores in each TMA to the average PPP across all six TMAs. Similarly, least squares linear regression was used to compare MPI for tissue cores in each TMA to the average MPI across all six TMAs. For each comparison, we calculated the slope (*m*) and coefficient of determination (*R*^*2*^). To assess concordance of MIBI-TOF with single-plex IHC, we used least squares linear regression to compare values for PPP attained by MIBI-TOF and IHC in adjacent serial tissue sections.

### Cell segmentation and phenotyping

To delineate the location of single cells in MIBI-TOF images, we performed cell segmentation using the pre-trained Mesmer convolutional neural network architecture.^24^ We used dsDNA as the nuclear marker and HLA class I and Na-K-ATPase as the membrane markers as input to the network. The output of Mesmer is the location of each cell in the image.

After cell segmentation, the next step of the analysis pipeline was to determine the phenotype of each individual cell. Pre-processed MIBI-TOF images were first Gaussian blurred using a standard deviation of 2 for the Gaussian kernel. Pixels were normalized by their total expression, such that the total expression of each pixel was equal to 1. A 99.9% normalization was applied for each marker. Pixels were clustered into 100 clusters using FlowSOM^25^ based on the expression of 12 phenotypic markers: CD3, CD4, CD8, CD11c, CD20, CD31, CD45, CD56, CD68, PanCK, Pax5, and Vimentin. The average expression of each of the 100 pixel clusters was found and the z-score for each marker across the 100 pixel clusters was computed. All z-scores were capped at 3, such that the maximum z-score was 3. Using these z-scored expression values, the 100 pixel clusters were meta-clustered using consensus hierarchical clustering into 12 meta-clusters. Next, by applying the segmentation masks that delineate the boundaries of all cells in the images, we counted the number of each of the 12 pixel clusters in each cell. This resulted in a pixel cluster by cell count table. These counts were then normalized by cell size. Using these frequency measurements as the feature vector, the cells were clustered using FlowSOM into 100 cell clusters. Similarly to the pixel clusters, the average expression of each of the 100 cell clusters was found and the z-score was computed. All z-scores were capped at 3, such that the maximum z-score was 3. Using these z-scored values, the 100 cell clusters were meta-clustered using consensus hierarchical clustering into 10 cell meta-clusters. Each of the cell meta-clusters was then manually annotated with its cell phenotype by assessing marker expression, resulting in a total of eight cell types: B cells, T cells, dendritic cells, macrophages, NK cells, fibroblasts, endothelial cells, and epithelial cells. To assess the concordance of cell phenotyping between serial sections of the same tissue core, we quantified the number of each cell type in each image, then calculated the average Spearman correlation of these frequencies between replicate images of each core.

## Results

### Experimental design for MIBI-TOF replicate and IHC concordance

To assess the reproducibility of MIBI as well as benchmark against single-plex chromogenic IHC, we took 13 serial sections from a TMA constructed for this study (see Methods) that included disease-free controls and multiple types of carcinomas, sarcomas, and central nervous system lesions (Figure 1). Here, every other slide was stained and processed for either MIBI-TOF analysis or single-plex chromogenic IHC for the indicated target, in addition to slides for H&E and unstained tissue control. To control for confounding factors due to batch effects resulting from day of tissue staining and imaging, the order that each TMA was stained and imaged were randomized with respect to serial sectioning order. Additionally, the order that each tissue core was acquired for each TMA was randomized as well (Table 1).

**Figure 1:**
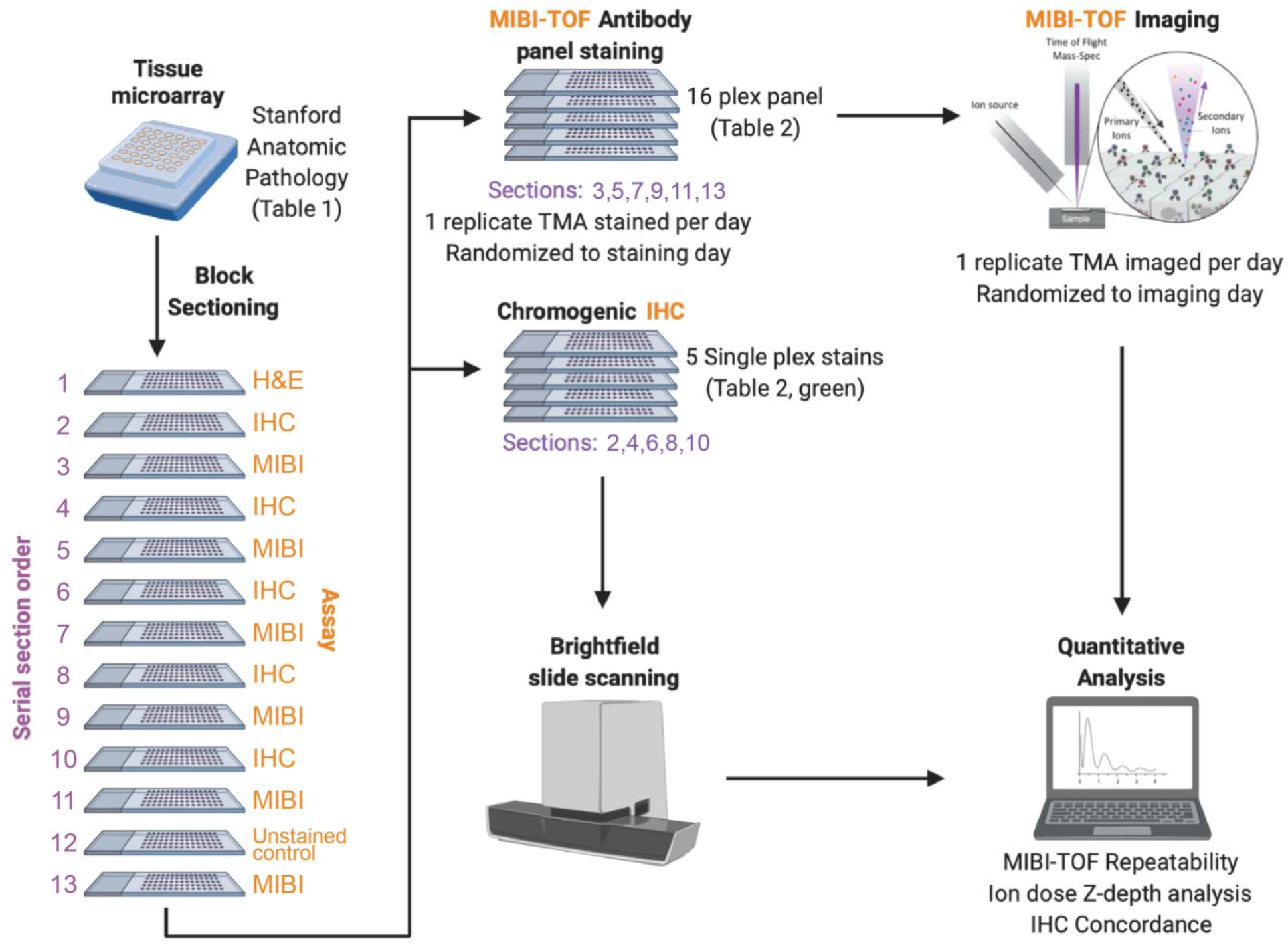
MIBI-TOF validation overview. A tissue microarray (TMA) was constructed using human FFPE tissue blocks, and the TMA was serially sectioned for analysis by single-plex chromogenic IHC and MIBI-TOF. The order that each serial section was stained and imaged was randomized. We then analyzed the concordance between IHC and MIBI-TOF, as well as assessed reproducibility of MIBI-TOF between serial sections.

### MIBI-TOF replicate concordance

For MIBI-TOF, 21 tissue cores in six replicate TMAs were stained with a 16-plex antibody panel and analyzed for a total of 126 MIBI-TOF images (Table 2). Instrument stability was assessed prior to data acquisition using a molybdenum foil standard, for which we know the expected range of ion counts. Primary ion current and mass resolution varied insignificantly over the course of the study (SD = 7% and 1.8%, respectively, Supplementary Figure 1A,B). Minor variations in ion detector sensitivity (SD = 22%) were within historical norms and were not found to impact subsequent quantitative comparisons (Supplementary Figure 1C). Fields in each tissue core were manually co-registered with respect to a slide scan of a reference H&E serial section, which showed high concordance of histological features (Supplementary Figure 2). High quality MIBI-TOF imaging data was obtained for all targets in the antibody staining panel in all six TMA slides. Manual evaluation of each marker demonstrated robust and specific staining to the target cells of interest in both normal and disease tissue cores (Figure 2).

**Figure 2:**
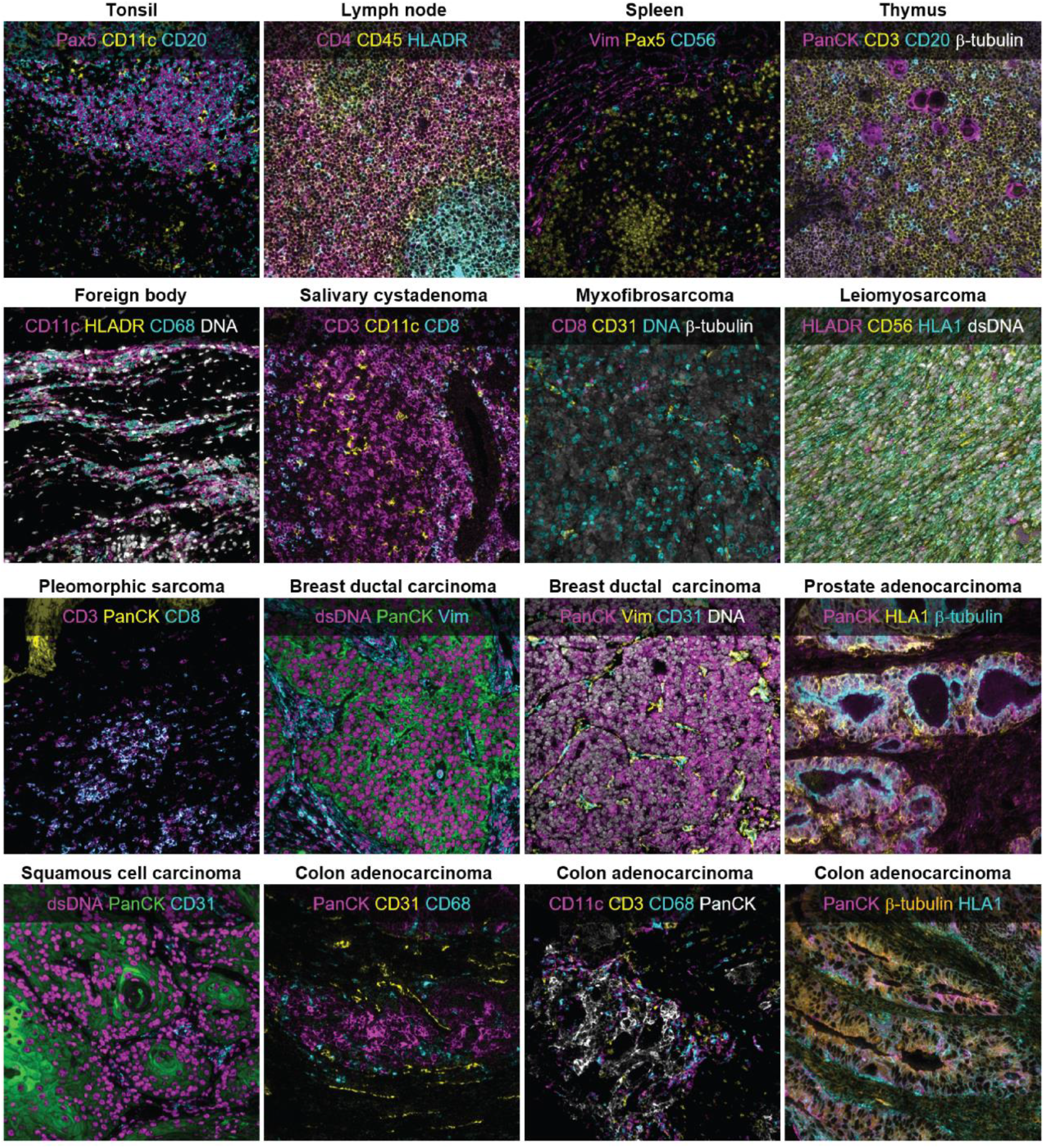
Representative MIBI-TOF imaging data. High-quality MIBI-TOF imaging data was attained for 16 markers for all replicate TMA slides. Representative images of the various tissue types in the TMA are shown here.

Assay reproducibility was assessed by comparing marker intensity and abundance in each TMA replicate. An example of co-registered fields-of-view (FOVs) from 6 serial sections of a representative tissue core is shown in Figure 3A, illustrating the high degree of reproducibility between the replicate sections. PPP values for CD3, a marker for T cells, and CD20, a marker for B cells, in the fields for this tissue core were 22.6% ± 2.5 and 20.6% ± 8.1 (mean ± SD), respectively. The significantly higher standard deviation observed with CD20 is attributable to section-to-section changes in the area occupied by two germinal centers (marked by expression of the B cell marker CD20) as sections were taken from the tissue core.

**Figure 3:**
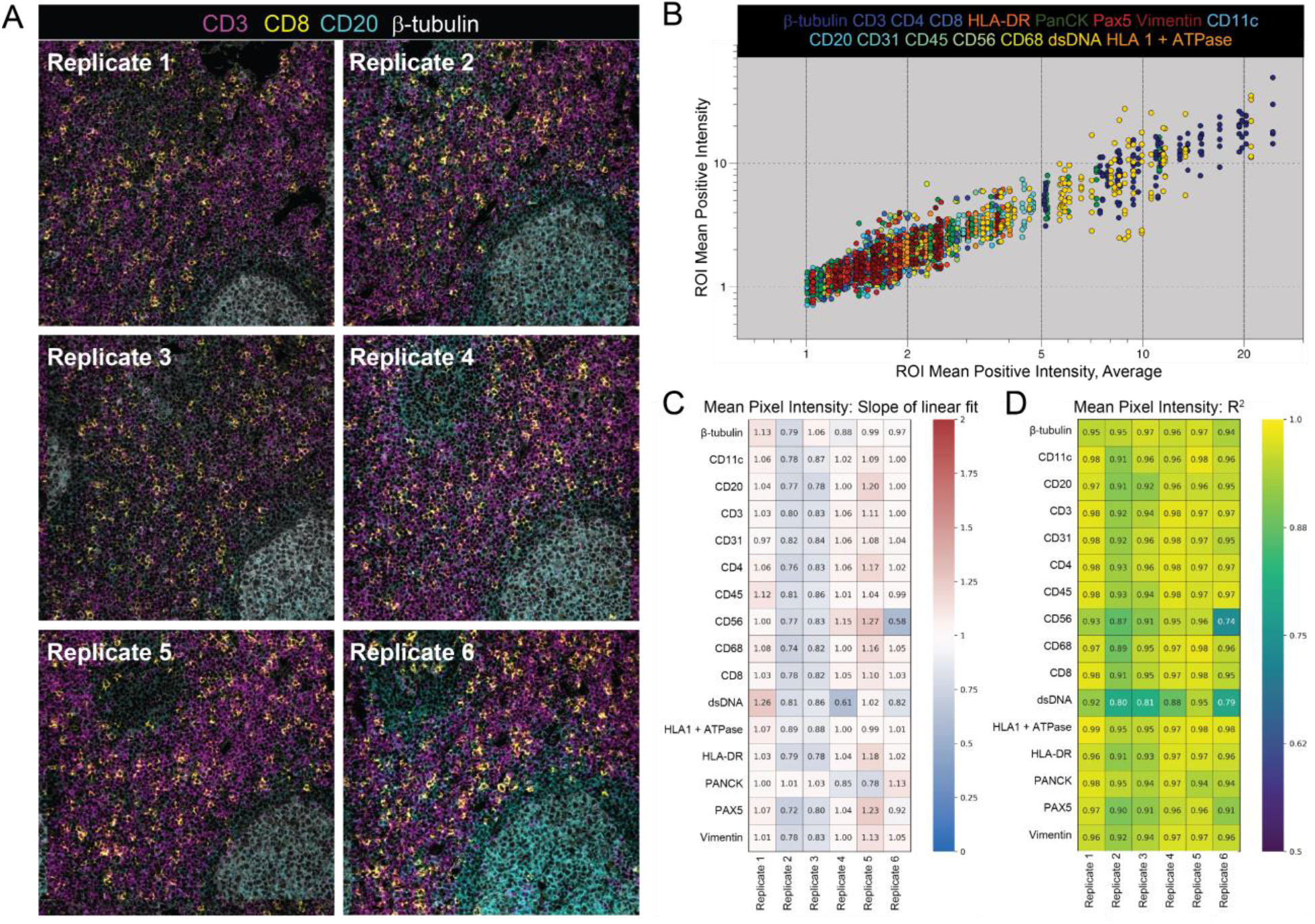
Concordance of Mean Pixel Intensity by MIBI-TOF. (A) Representative MIBI-TOF replicate images from serial sections of the same TMA core of lymph node tissue. (B) Plot of Mean Positive Intensity (MPI) of each FOV vs. the average MPI of all FOVs of the same TMA core. Each color represents a different marker. We performed least squares linear regression of the MPI of all six replicates of each TMA core vs the average MPI of the core and found the slope *m* (C) and coefficient of determination *R*^*2*^ (D) for all 16 markers.

Linear regression of normalized values for marker intensity (MPI) and marker frequency (PPP) for each tissue core in each TMA replicate (see Methods) were used to determine the degree of concordance with respect to the average across all six replicates. For MPI, which quantifies the average marker intensity in positive pixels, the resultant slope (*m*) and coefficient of determination (*R*^*2*^) values for all markers in all six replicates were close to 1, with a *m* of 0.96 ±

0.14 and a *R*^*2*^ of 0.94 ± 0.04 (mean ± SD, Figure 3B-D). MPI captures marker intensity, thus *m* and *R*^*2*^ values close to 1 indicate that when a marker is present, the brightness of the marker is highly concordant across replicates. Similarly, for PPP, which quantifies the total number of pixels that are positive for a given marker, *m* and *R*^*2*^ in all replicates and markers were also close to 1, with a *m* of 0.98 ± 0.15 and a *R*^*2*^ of 0.95 ± 0.04 (mean ± SD, Supplementary Figure 3). PPP *m* and *R*^*2*^ values close to 1 indicate that the number of positive pixels across replicates of the same core is consistent. Taken together, these two metrics demonstrate that MIBI-TOF images of serial sections of the same tissue core that were randomized for staining and imaging day were highly concordant, lending evidence for the reproducible nature of MIBI-TOF.

### Reproducibility of cell immunophenotypic assignments by MIBI-TOF

An important step in the analysis of high-dimensional imaging data is the identification of single cell lineages by immunophenotype in the images. Thus, we next assessed the reproducibility of single-cell annotations across the replicate images of each tissue core. Our lab has recently developed a cell segmentation algorithm that can accurately identify the location of cells in solid tissue without the need for manual fine-tuning or user adjustment.^24^ We used this method to delineate the location of single cells in the images, then used a pixel-clustering based approach to classify the single cells into eight cell lineages: B cells, T cells, dendritic cells, macrophages, NK cells, fibroblasts, endothelial cells, and epithelial cells (Figure 4A, see Methods).^17^ After classifying the cells in each of the images, we quantified the number of each cell type in each image and assessed concordance between replicates by calculating the average

**Figure 4:**
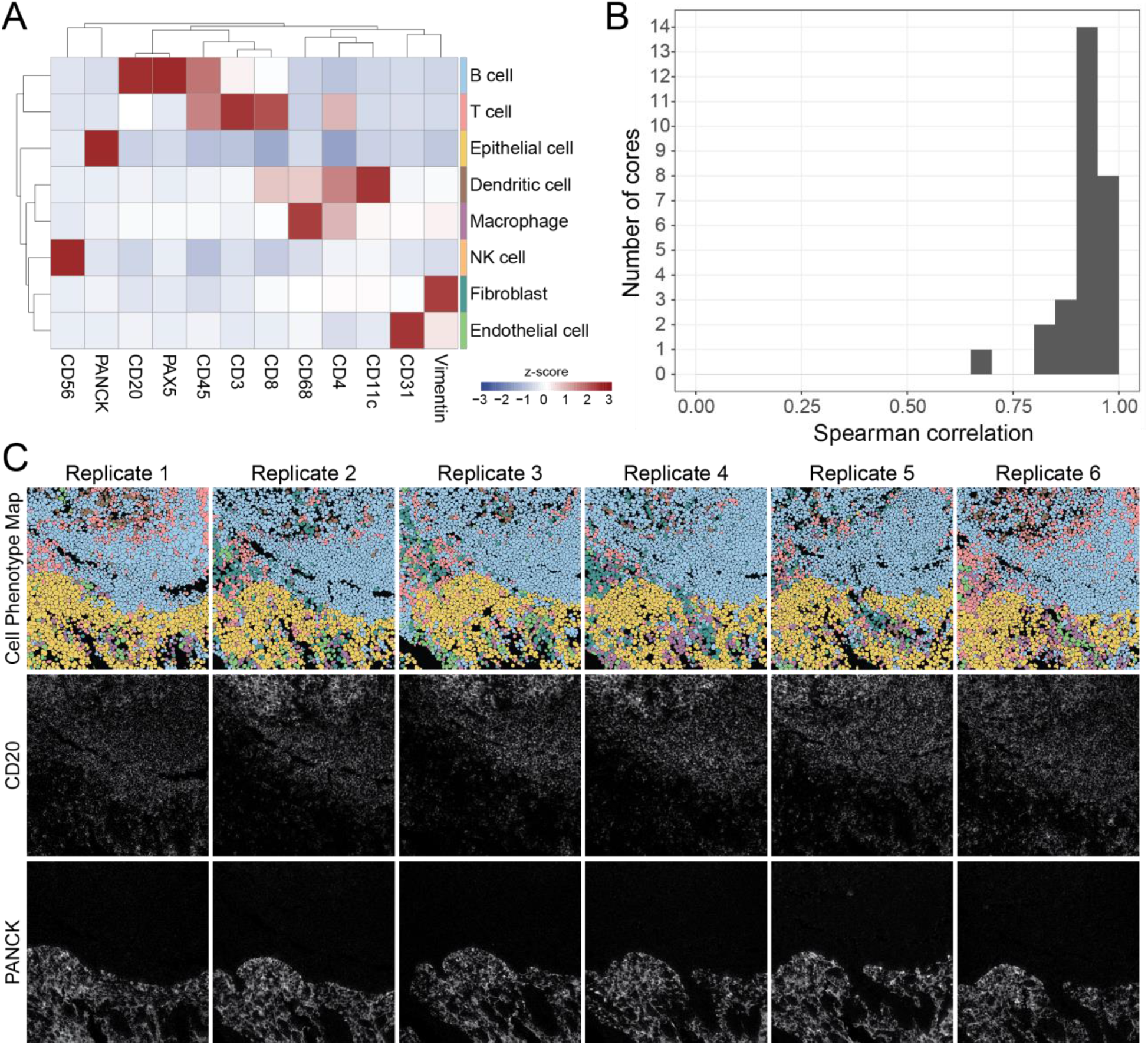
Reproducibility of cell phenotyping by MIBI-TOF. (A) Cells were assigned to a phenotype using an unsupervised clustering approach and manually annotated into eight major cell types. Expression values were z-scored for each marker. High z-scores are an indicator of high marker specificity. (B) The Spearman correlation between all serial sections of each TMA core using the frequency of cell types in each FOV. (C) Representative images of six serial sections of the same TMA core of tonsil tissue. Top row: Cell phenotype map colored according to the eight cell types shown in (A). Middle and bottom rows: MIBI-TOF images showing CD20 and PanCK expression.

Spearman correlation of these frequencies between the replicate images of each tissue core (Figure 4B). We expected variation in these Spearman correlation values, since the replicate sections of the same tissue core are serial sections of 4 μm each (with additional serial sections taken for IHC concordance analysis). In total, 13 serial sections were obtained for each tissue for MIBI-TOF or IHC (see Methods). Therefore, we would expect true biological differences when comparing serial sections up to 48 μm apart. When comparing these replicate MIBI-TOF images, we observed that broad histological features were conserved, such as germinal centers or vessels (Figure 3A, 4C, Supplementary Figure 4). The average Spearman correlation was high, 0.92 ± 0.06 (mean ± SD), demonstrating that cell phenotyping, a necessary step in high-dimensional image analysis, was reproducible between MIBI-TOF images of the same tissue core.

### Concordance of MIBI-TOF with single-plex chromogenic IHC

Chromogenic IHC is the standard modality for disease prognosis and therapeutic selection in the vast majority of solid tumors. Therefore, we assessed concordance to determine if multiplexed imaging by MIBI-TOF can quantitatively recapitulate single-plex immunoassays. Similar to the analyses described above that were used to compare MIBI-TOF data across replicates, the frequency of marker positive pixels in MIBI-TOF images co-registered with single-plex chromogenic IHC stains were compared by linear regression across all 21 tissue cores. Because we alternated recuts for MIBI-TOF and IHC, we could compare MIBI-TOF and IHC on adjacent serial sections. Single-plex chromogenic IHC images were acquired for five targets: CD3, CD8, Pax5, PanCK, and CD68. Marker positivity by single-plex IHC for CD3, CD8, Pax5, PanCK, and CD68 in adjacent serial sections were compared with the respective values found by MIBI-TOF using linear regression (Figure 5, Supplementary Figure 5). For all five targets, concordance of MIBI-TOF and IHC was high, with an *R*^*2*^ of 0.85 ± 0.08 (mean ± SD, all *R*^*2*^ > 0.7), demonstrating that MIBI-TOF data is consistent with single-plex chromogenic IHC data.

**Figure 5:**
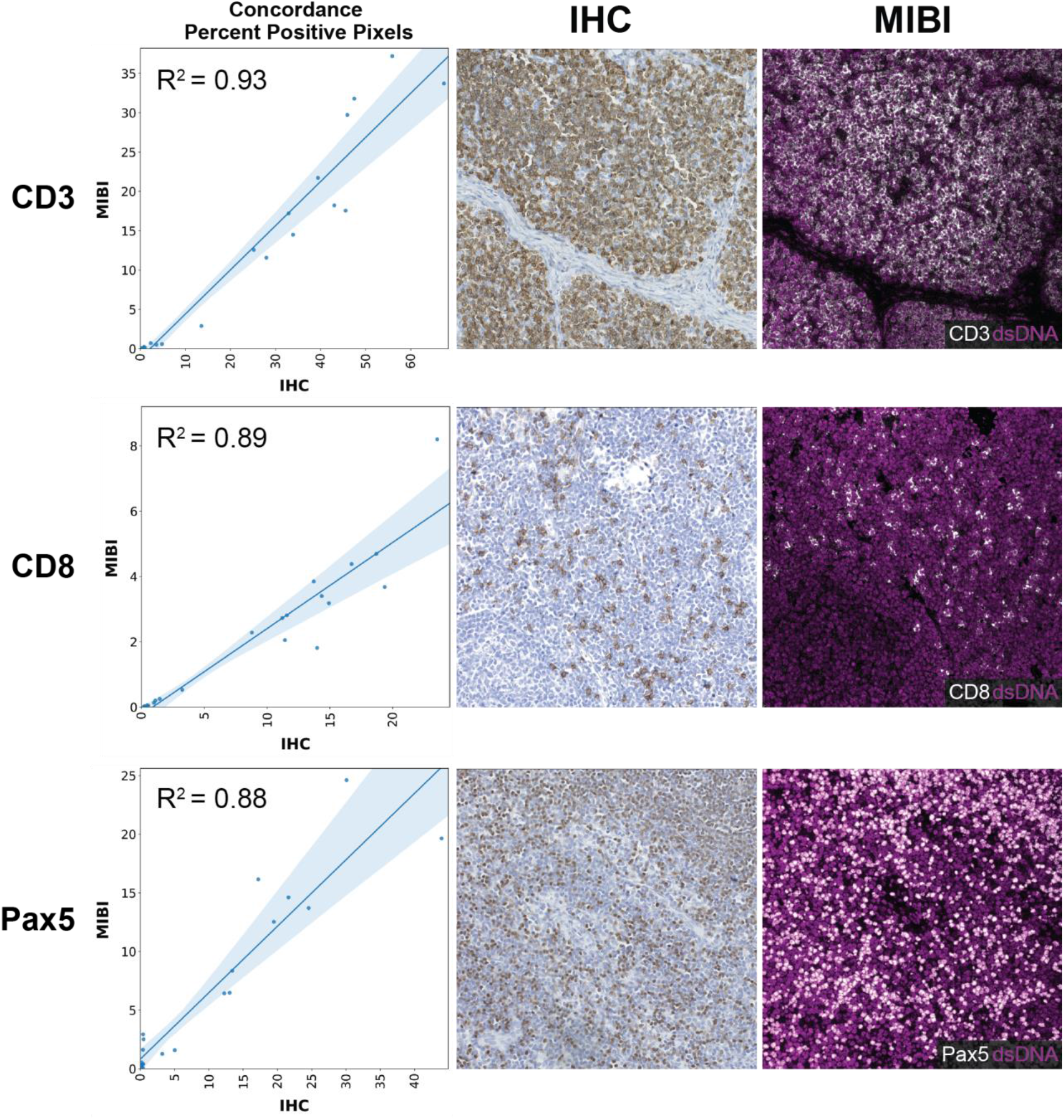
Concordance of MIBI-TOF with single-plex chromogenic IHC. Representative images of the comparison of MIBI-TOF images co-registered with single-plex chromogenic IHC stains. Each data point represents the PPP by MIBI-TOF and IHC for a single tissue core. Shaded area represents 95% confidence interval. Additional markers are shown in Supplementary Figure 5.

## Discussion

In this study, we assessed the reproducibility of MIBI-TOF by evaluating assay concordance on six serial sections from a TMA, including disease-free tonsil, lymph node, thymus, and spleen, in addition to multiple types of carcinomas, sarcomas, and central nervous system lesions. Importantly, each serial section was stained independently on six separate days with a 16-marker staining panel and subsequently analyzed on six different days. 21 tissue cores were analyzed in each serial section for a total of 126 MIBI-TOF images. The staining intensity and frequency of positive pixels for each serial section (MPI and PPP, respectively) were compared against average values across all runs using least squares linear regression. In all comparisons, concordance between serial sections of the same tissue core was high, with *m* of 0.96 ± 0.14 and *R*^*2*^ of 0.94 ± 0.04 when assessing MPI and *m* of 0.98 ± 0.15 and *R*^*2*^ of 0.95 ± 0.04 when assessing PPP (mean ± SD). The staining pattern of each marker was manually inspected to confirm, where relevant, appropriate subcellular localization and histologic distribution. For example, CD20 staining in positive regions of tonsil and lymph node in all replicates was verified to be membrane-localized and enriched in follicles. Furthermore, we assessed the reproducibility of cell phenotyping across replicates and found that across replicate sections, the location and lineage of cells identified using our data analysis pipeline were highly concordant, with an average Spearman correlation of cell type frequencies between replicates of 0.92 ± 0.06 (mean ± SD). Importantly, the data analysis pipeline used to go from MIBI-TOF images to cell phenotypes was nearly fully automated. Cell segmentation and clustering were fully automated with no user input and only minimal user intervention was needed when labelling cell clusters with cell annotations, demonstrating that the biological annotation of MIBI-TOF images is robust without the need for significant researcher intervention. In addition, we assessed concordance of PPP by MIBI-TOF for five markers with single-plex IHC and found that MIBI-TOF and single-plex IHC were highly concordant with an average *R*^*2*^ of 0.85. Taken together, the combination of consistent staining, imaging, and feature extraction illustrates the highly reproducible nature of MIBI-TOF.

One limitation of validation studies involving serial tissue sections is that exact biological replicates cannot be obtained since each tissue section is somewhat different and is not a homogenized bulk sample. In this study, since the tissue sections are 4 μm each and a total of 13 serial sections were acquired, the first and last tissue section were separated axially by 48 μm. Also, when balancing assay replication versus concordance with an orthogonal assay, like single-plex IHC here, one needs to prioritize what physically adjacent slides are used for. In this case, we selected physically adjacent slides for benchmarking single-plex IHC since any one marker’s distribution would be heavily dependent on cell composition. Lymphocytes that are approximately ∼10 μm would be best preserved in adjacent sections. Although there will be true biological differences in this volume of tissue, this study represents the next best situation, in which we compared adjacent serial sections as a proxy for true replicates. Furthermore, this was a single-site study, in which all staining and imaging was performed in our laboratory. We are currently planning a multi-site study to assess inter-institutional differences in instrument performance and tissue staining. Still, we believe the overall metrics for benchmarking outlined here will be sufficient to assess these future efforts.

This study was performed as a part of the Cancer Immune Monitoring and Analysis Centers Cancer Immunology Data Commons (CIMAC-CIDC) network^22^, which is a NCI Cancer Moonshots initiative to provide the technology and expertise for immunotherapy clinical trials. Multiplexed tissue imaging is an integral assay for fully characterizing the tumor immune microenvironment, which requires the simultaneous profiling of multiple tumor and immune cell types. MIBI-TOF can routinely provide quantitative, multiplexed imaging data that is back compatible with archival FFPE tissue and conventional anatomic pathology workflows. Technical innovations in reagents and instrumentation will further increase the throughput and multiplexing ability of MIBI-TOF. In the future, we envision that MIBI-TOF will not only be used for reproducible basic science research, but also enable the adoption of quantitative spatial signatures in the clinic for more accurate diagnosis and therapeutic selection. At the same time, as efforts to develop other IHC-centric spatial imaging technologies evolve, the analytical framework and approach laid out in this study should serve as a guide for their assessment.

## Data Availability

The datasets generated and/or analyzed during the current study are available in Zenodo: 10.5281/zenodo.5542727.

## Acknowledgements

The authors thank Sushama Varma and Matt van de Rijn for creating the tissue microarray for this study.

## Conflict of Interest

M.A. and S.C.B. are inventors on patents related to MIBI technology. M.A. and S.C.B. are consultants, board members and shareholders in Ionpath Inc.

## Author Contributions

M.B. performed experiments and imaging. A.K., A.K., and C.C.L. performed data analysis. C.C.L. and M.A. wrote the manuscript. S.C.B., R.K., and S.M.H. reviewed the manuscript. S.C.B. and M.A. supervised the work.

## Funding

This work was supported by DOD EOH W81XWH2110143, 1-DP5-864 OD019822, 1R01AG056287, 1R01AG057915, R01AG068279, 1UH3CA246633, 1U24CA224309, and the Bill and Melinda Gates Foundation.

## Notes

https://doi.org/10.5281/zenodo.5542727

