## Supplementary Figures and Tables for "Reproducible, high-dimensional imaging in archival human tissue by Multiplexed Ion Beam Imaging by Time-of-Flight (MIBI-TOF)"

**Supplementary Figure 1: MIBI-TOF instrument stability**

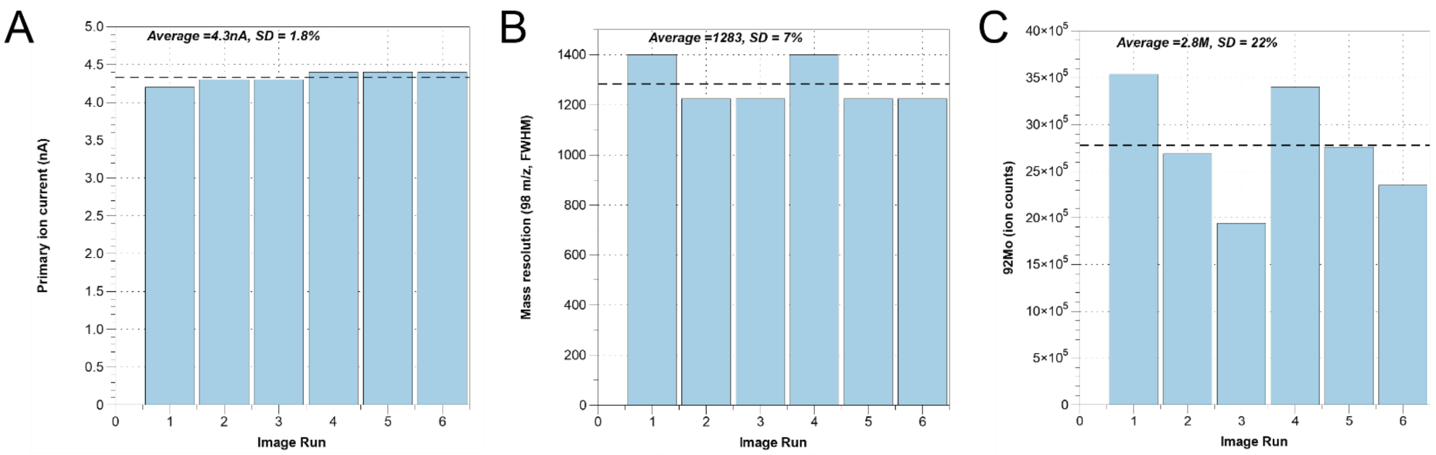


**Figure S1.** MIBI-TOF instrument stability was assessed prior to data acquisition using a molybdenum foil standard. (A) Primary ion current, (B) mass resolution, and (C) ion detector sensitivity were assessed and did not vary significantly over the course of the study.

**Supplementary Figure 2: Co-registration with brightfield microscopy**

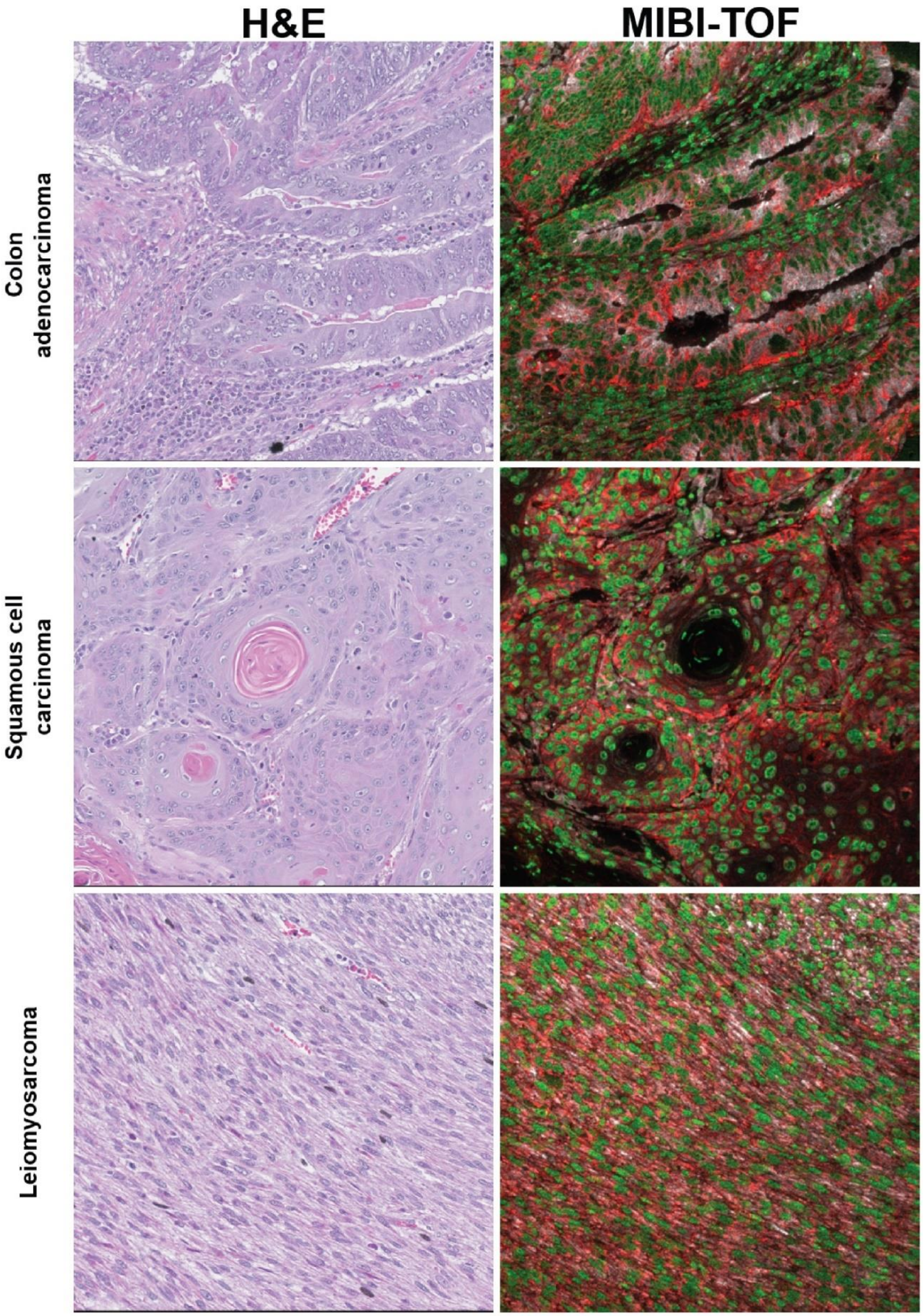


**Figure S2.** Representative co-registered H&E and MIBI-TOF images of 3 tissue cores. Green: dsDNA, red: HLA1 + ATPase.

#### Supplementary Figure 3: Concordance of Percent Positive Pixels by MIBI-TOF

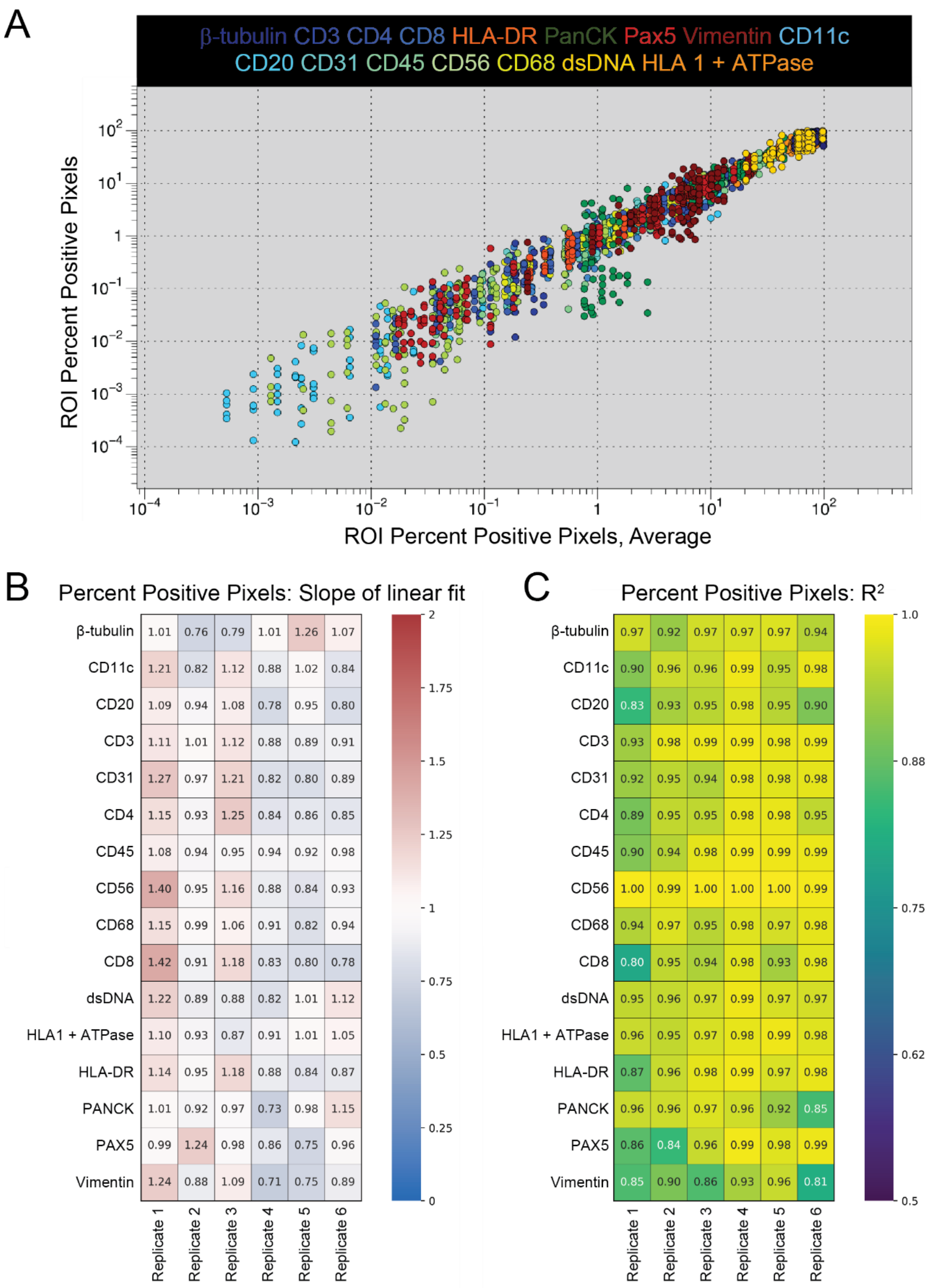


**Figure S3.** (A) Plot of Percent Positive Pixels (PPP) of each FOV vs. the average PPP of all FOVs of the same TMA core. Each color represents a different marker. We performed least squares linear regression of the PPP of all six replicates of each TMA core vs the average PPP of the core and found the slope *m* (B) and coefficient of determination $R^2$ (C) for all 16 markers.

**Supplementary Figure 4: Reproducibility of cell phenotyping by MIBI-TOF**

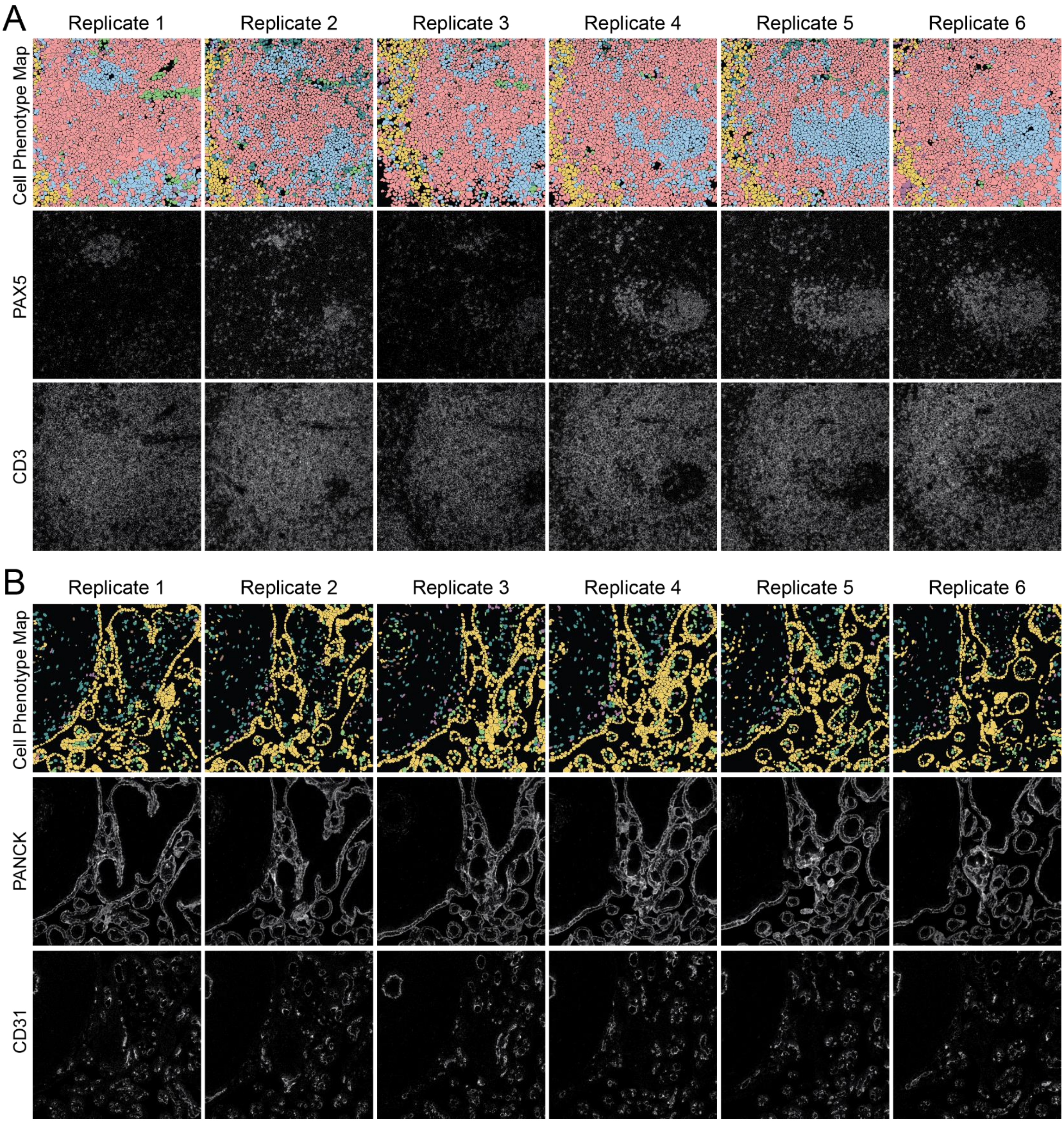


**Figure S4.** Additional examples showing cell type assignment reproducibility across six serial sections of the same TMA core of tonsil tissue (A) and placenta (B). In both (A) and (B), top row: cell phenotype map colored according to the eight cell types shown in Figure 4A; middle and bottom rows: single-marker MIBI-TOF images.

**Supplementary Figure 5: Concordance of MIBI-TOF with single-plex chromogenic IHC**

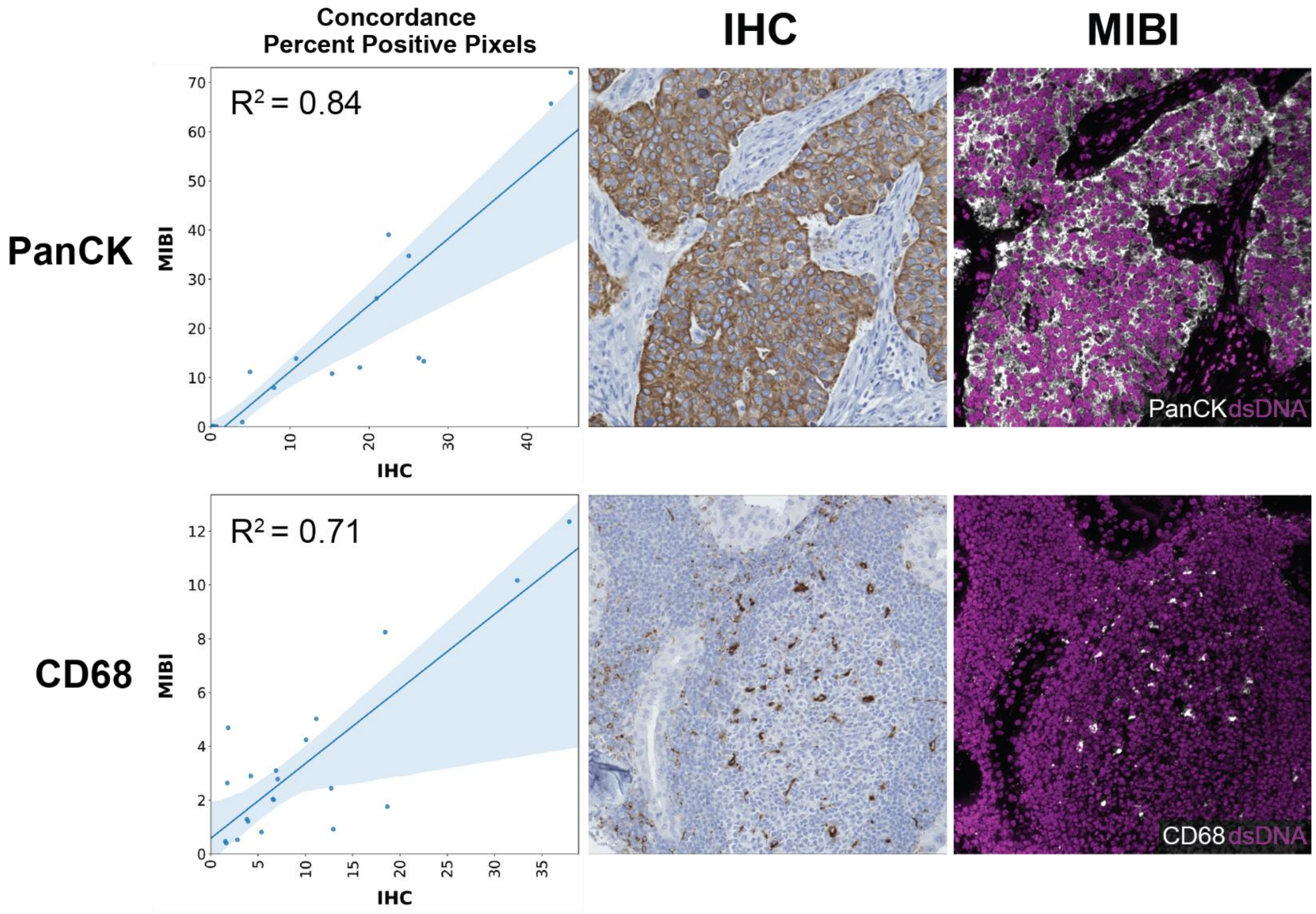


**Figure S5.** Representative images of the comparison of MIBI-TOF images co-registered with single-plex chromogenic IHC stains. Each data point represents the PPP by MIBI-TOF and IHC for a single tissue core. Shaded area represents 95% confidence interval.

### Supplementary Table 1: MIBI-TOF background correction parameters

| Channel | Mass Window | | Slide Background | | | Isobaric Correction | | | | | |
|---|---|---|---|---|---|---|---|---|---|---|---|
| | Start [$m_a$] | Stop [$m_a$] | $^{138}$Ba [counts] | $^{181}$Ta [counts] | $^{197}$Au [counts] | dsDNA ($^{89}$Y) | CD4 ($^{143}$Nd) | CD56 ($^{151}$Eu) | CD68 ($^{156}$Gd) | CD3 ($^{159}$Tb) | HLA DR ($^{172}$Yb) |
| dsDNA ($^{89}$Y) | 88.70 | 89.00 | | 20 | 50 | | | | | | |
| beta-tubulin ($^{113}$In) | 112.70 | 113.00 | | 20 | 50 | 3.30% | | | | | |
| CD4 ($^{143}$Nd) | 142.70 | 143.00 | | 20 | 50 | | | | | | |
| CD11c ($^{144}$Nd) | 143.70 | 144.00 | 15 | 20 | 50 | | 2.00% | | | | |
| CD56 ($^{151}$Eu) | 150.70 | 151.00 | 15 | 20 | 50 | | | | | | |
| CD31 ($^{152}$Sm) | 151.70 | 152.00 | 15 | 20 | 50 | | | | | | |
| CD68 ($^{156}$Gd) | 155.70 | 156.00 | 15 | 20 | 50 | | | | | | |
| CD8 ($^{158}$Gd) | 157.70 | 158.00 | | 20 | 50 | | | | 0.96% | | |
| CD3 ($^{159}$Tb) | 158.70 | 159.00 | | 20 | 50 | | | | | | |
| Vimentin ($^{163}$Dy) | 162.70 | 163.00 | 15 | 20 | 50 | | | | | | |
| PAX5 ($^{165}$Ho) | 164.70 | 165.00 | 15 | 20 | 50 | | | | | | |
| CD20 ($^{167}$Er) | 166.70 | 167.00 | | 20 | 50 | | 1.50% | | | | |
| HLA DR ($^{172}$Yb) | 171.70 | 172.00 | | 20 | 50 | | | | | | |
| PANCK ($^{173}$Yb) | 172.70 | 173.00 | | 20 | 50 | | | | 5.00% | 1.00% | 5.20% |
| CD45 ($^{175}$Lu) | 174.70 | 175.00 | | 20 | 50 | | | 1.00% | | | |
| HLA class 1 A, B, and C Na-K-ATPase alpha ($^{176}$Yb) | 175.70 | 176.00 | | 20 | 50 | | | | | | |

#### Supplementary Table 2: MIBI-TOF noise filtering parameters

| Channel | Aggregates diameter [um] | Two intensity levels | Initial noise estimation | Reachability cut-off | | Reachability mask |
|---|---|---|---|---|---|---|
| | | | | Slope | Intercept | |
| dsDNA ($^{89}$Y) | | | Reachability | -0.16 | 1.16 | |
| beta-tubulin ($^{113}$In) | | TRUE | Conjunction | 0.00 | 1.25 | |
| CD4 ($^{143}$Nd) | 2.5 | | Max | -0.44 | 1.57 | TRUE |
| CD11c ($^{144}$Nd) | 2.5 | | Max | -0.29 | 1.43 | TRUE |
| CD56 ($^{151}$Eu) | 2.5 | | Reachability | -0.25 | 1.25 | TRUE |
| CD31 ($^{152}$Sm) | | | Reachability | -0.52 | 0.90 | TRUE |
| CD68 ($^{156}$Gd) | | | Max | -0.37 | 1.21 | TRUE |
| CD8 ($^{158}$Gd) | | | Reachability | -0.34 | 1.41 | TRUE |
| CD3 ($^{159}$Tb) | 2.5 | | Reachability | -0.32 | 1.50 | TRUE |
| Vimentin ($^{163}$Dy) | | | Reachability | -0.24 | 1.37 | |
| PAX5 ($^{165}$Ho) | 2.5 | | Max | -0.27 | 1.41 | TRUE |
| CD20 ($^{167}$Er) | 2.5 | | Connectivity | -0.33 | 1.39 | TRUE |
| HLA DR ($^{172}$Yb) | | | Reachability | -0.41 | 1.27 | TRUE |
| PANCK ($^{173}$Yb) | | TRUE | Reachability | -0.30 | 1.60 | |
| CD45 ($^{175}$Lu) | 2.5 | | Reachability | -0.22 | 1.53 | TRUE |
| HLA class 1 A, B, and C Na-K-ATPase alpha ($^{176}$Yb) | | TRUE | Conjunction | -0.26 | 1.78 | |

#### Supplementary Table 3: IHC thresholds

| Channel | DAB Threshold | Hematoxylin Threshold |
|---|---|---|
| CD68 | 0.130 | 0.210 |
| CD8 | 0.126 | 0.210 |
| CD3 | 0.177 | 0.168 |
| PAX5 | 0.169 | 0.207 |
| PANCK | 0.110 | 0.262 |
